# Changing environments reveal innovative genetic variation in children’s cortisol responses

**DOI:** 10.1101/856658

**Authors:** Laurel Raffington, Margherita Malanchini, Andrew D. Grotzinger, James W. Madole, Laura E. Engelhardt, Aditi Sabhlok, Cherry Youn, Megan W. Patterson, K. Paige Harden, Elliot M. Tucker-Drob

**Affiliations:** Department of Psychology, University of Texas at Austin, United States; Department of Biological and Experimental Psychology, Queen Mary University of London

**Keywords:** Gene-environment interaction, reaction norms, stress, cortisol

## Abstract

Genetic associations with biopsychosocial phenotypes are often interpreted as evidence that the genome codes for fixed end-states. Instead, a given genotype might regulate a dynamic range of phenotypes in response to environmental change. We collected hair cortisol (*n* = 1,104), salivary cortisol in reaction to an in-laboratory stressor (*n* = 537), and diurnal salivary cortisol (*n* = 488) from twins aged 8-15 years in the Texas Twin Project. Baseline genetic variation in both salivary and hair cortisol was not simply magnified after stressor exposure or after waking. Rather, novel genetic influences on cortisol arose over time. Thus, environmental change can reveal genetic variation that would not otherwise be observed in static cortisol levels. These findings are in line with the notion that the genome regulates individuals’ reactions to the environment that differ across environments.

---

Genetic effects on biopsychosocial phenomena are often interpreted as evidence that the genome predisposes an individual organism to a fixed phenotypic end-state (Burt, 1966). A competing view contends that genetic variation modulates how organisms react to the environment (Gottlieb, 2007). Such genetic response patterns have been called reaction norms (Dobzhansky, 1955; Woltereck, 1909 cf Platt & Sanislow, 1988) or reaction ranges (Gottesman, 1963; *see* Griffiths & Tabery, 2008 *for a historical discussion of the two concepts*). A key theoretical debate has been whether mutable environments (a) change mean levels of a phenotype without altering the genetic basis of individual differences therein (*e.g.*, as seen in Capron & Duyme, 1989); (b) overwhelm genetic influences; (c) amplify pre-existing genetic differences; or (d) evoke different genetic variation (*i.e.*, innovative genetic variation), such that the relative ordering of phenotypes in novel environments is unpredictable from ordinarily observed or pre-existing individual differences (*e.g.*, Gottlieb, 2007; Gupta & Lewontin, 1982). Empirical examinations of these research questions have primarily been confined to non-human organisms, such as fruit flies (Gupta & Lewontin, 1982) and mice (Cooper & Zubek, 1958). Here, we examine the same theoretical question in a behaviorally-relevant human phenotype.

Documenting how genetic variation relates to responses to environments (Plomin, DeFries, & Loehlin, 1977) is hampered by three primary challenges. First, most complex human phenotypes are affected by a very large number of genetic variants with individually small effect sizes (Chabris, Lee, Cesarini, Benjamin, & Laibson, 2015). Thus, previous studies of interactions between individual genetic variants and environmental exposures have largely failed to replicate (Border et al., 2019; Halldorsdottir & Binder, 2017). Second, most studies of differences in genetic influence by environment have examined different individuals in different environments (*e.g.*, Tucker-Drob & Bates, 2016) without examining how genetic effects differ as environments change for the same group of individuals. Third, most studies of gene × environment interaction have measured naturally-occuring enviornmental variation without attempting to manipulate enviornments (Schmitz & Conley, 2017). Therefore, it is difficult to discern whether enviornmental measures represent exogenous influences on the individuals versus endogenous factors that are themselves correlated with genetic variation, or with other factors relevant for the outcomes under study (Conley, 2011).

Here, we overcome these past challenges in a study of the stress-sensitive hormone cortisol. Cortisol secretion is an environmentally-responsive, psychologically-relevant biomarker that is well-suited for investigating the intersection of genetic variation with environmental change. Measures of cortisol secretion are commonly investigated in relation to changing environments, even over relatively short timescales. For example, exogenously-manipulated stress increases salivary cortisol output within minutes (Hellhammer, 2011). Similarly, salivary cortisol increases drastically upon awakening (cortisol awakening response) and decreases over the rest of the day (diurnal slope; Miller et al., 2016). Cumulative cortisol levels, as measured in hair-based assays, have been found to be associated with chronic, more temporally stable forms of environmental variation (Dajani, Hadfield, van Uum, Greff, & Panter-Brick, 2018). Both salivary and hair cortisol measures have been linked with a range psychological and behavioral outcomes (e.g., Bäumler et al., 2014; Shalev et al., 2019; White et al., 2017). More generally, cortisol secretion is considered relevant for a variety of behavioral and health outcomes (Lupien, McEwen, Gunnar, & Heim, 2009).

Importantly, although cortisol secretion is sometimes primarily interpreted as a biomarker of environmental exposure, it is also heritable (Rietschel et al., 2017; Tucker-Drob et al., 2017). The majority of previous studies investigating genetic variation of cortisol secretion have examined static cortisol levels rather than changes in cortisol over time (Rietschel et al., 2017; Tucker-Drob et al., 2017). The few studies that have examined the contribution of genetic variation to cortisol reactions or diurnal change have reported sizable heritability estimates (Federenko, Nagamine, Hellhammer, Wadhwa, & Wüst, 2004; Ouellet-Morin et al., 2016; Van Hulle, Shirtcliff, Lemery-Chalfant, & Goldsmith, 2012). It remains unknown whether heritable contributions to cortisol response simply reflect magnification of standing genetic contributions to baseline variation in cortisol, or whether there are novel contributions of genetic factors not evident prior to stressor onset.

In this study, we apply behavior genetic methods to data from up to 1,104 individuals from a population-based sample of grade school twins. We estimate genetic variation in hair-based chronic levels, baseline salivary cortisol levels, and in changes in salivary cortisol both across the day and in response to a standardized in-laboratory stressor. We also estimate genetic correlations across cortisol levels and responses to answer the following three research questions:

1. Is there distinct genetic variation relevant to cortisol reactions to acute stress (*i.e.* a novel in-laboratory stressor) and cortisol secretion across the day (*i.e.* diurnal cortisol fluctuation)?
2. Do changes in cortisol amplify standing genetic variation in baseline levels of cortisol, or do they reveal innovative genetic variation?
3. Are cortisol reactions to an acutely stressful environment and cortisol secretion across the day regulated by the same genetic factors?

## Method

### Participants

Participants were members of the Texas Twin Project, an ongoing longitudinal, population-based study of twins and multiples living in the Austin metropolitan area (Harden, Tucker-Drob, & Tackett, 2013). Families were recruited from public school rosters. Twin pairs included in the current analyses had at least one measure of cortisol, had no hormone-disrupting disorder (*n* = 12), and had not taken steroid-based medication regularly in the past six months (*n* = 19; total exclusion *n* = 27). The final sample (*N* = 1,104 unique individuals, 53% female) consisted of 150 monozygotic and 304 dizygotic twin pairs from 454 families (see Supplement for zygosity classification). Participants ranged in age from 8 to 15 years (*M* = 11.01, *SD* = 1.81). 204 families contributed data at more than one wave of data collection (mainly hair cortisol, up to three waves). Sample size varied depending on the cortisol collection modality (see Table S1). Twin pairs identified as being best described as White (62.11%), Latinx (13.88%), Latinx-White (8.59%), African-American (3.52%), Asian (4.40%), or other multiracial/multiethnic combinations (7.50%). The University of Texas at Austin Institutional Review board granted ethical approval.

### Measures

#### Hair cortisol

1,078 unique participants contributed at least one measure of hair cortisol. Several participants contributed multiple samples of this ongoing longitudinal study, resulting in 1,338 hair cortisol samples including repeated measures. Procedures for accounting for the nesting of data within individuals are described in the Analyses section below. Research assistants collected a hair sample approximately 3 mm wide and 3 cm long from the posterior vertex of the scalp; this served as a marker for average cortisol secretion over the most recent 3-month period. See Supplemental Methods for more collection details. Hair cortisol values were residualized for assay batch (separately from salivary cortisol) and log-transformed.

#### Cortisol reactions to acute stress

Cortisol reactions to stress were measured using the Trier Social Stress Test (Kirschbaum, Pirke, & Hellhammer, 1993) adapted to children (TSST-C; Buske-Kirschbaum et al., 1997). Approximately 30 min after their arrival at the lab, participants were instructed to prepare a short story to be presented in front of two judges. After a 5 min preparation period, they then presented the story (5 min) and were asked to orally calculate mental arithmetic problems in front of the judges (5 min). Four salivary samples indexed participants’ cortisol before, during, and following the TSST-C protocol: (1) shortly upon arrival to the lab and at least 30 minutes before the TSST–C, (2) 20 minutes after the start of the TSST– C, (3) 20 minutes after the completion of sample 2, and (4) 20 minutes after the completion of sample 3. See Supplemental Methods for more details on data collection.

537 unique participants contributed at least one measure of cortisol secretion in reaction to the TSST-C (Table S1). Salivary cortisol values were residualized for batch (the year the lab assayed the sample) and log-transformed to correct for positive skew.

#### Diurnal cortisol secretion

Saliva collection kits were provided for four consecutive days with an additional fifth kit in case of sampling problems, which 22% of participants completed in spite of not having experienced any sampling problems. Samples were taken at home with the help of parents three times a day: immediately upon waking, 30 min after waking, and right before bedtime. See Supplemental Methods for more details on data collection and processing.

488 unique participants contributed at least one measure of diurnal cortisol secretion. Several participants contributed diurnal data over multiple collection waves, resulting in 574 unique sets of sampling days including repeated measures (see Table S1 for descriptive statistics). Procedures for accounting for the nesting of data within individuals are described in the Analyses section below. Salivary cortisol values were residualized for analytic batch (together with stress cortisol measures), as well as non-steroid medication use on the day of sampling, dairy consumption, and waking time on that day. Residualized cortisol values were log-transformed to correct for positive skew.

### Analyses

#### Phenotypic stress reaction and diurnal cortisol models

Following the modeling approach of Malanchini et al. (2019), we applied multilevel piecewise latent growth models to characterize the change in salivary cortisol at the intra- and inter-individual levels. Within this two-level growth model framework, Level 1 represented within-person variation in the cortisol trajectory, and Level 2 denoted between-person variation after controlling for the effect of intra-individual variability. Time was scaled in hours (such that, e.g. the mean diurnal slope can be interpreted as a rate of change in transformed cortisol residuals per hour).

At Level 1, we specified three latent factors to characterize cortisol levels surrounding the acute stressor: (1) an intercept that reflects pre-stress baseline cortisol levels, (2) a latent response slope capturing the rise in cortisol following stress, and (3) a latent recovery slope representing the decline in cortisol following the response. The model estimated the rise in cortisol response prior to a specified turning point and an independent recovery slope following the turning point, which was found to be optimal 25 minutes from the start of the TSST-C in the present sample (Malanchini et al., 2019). Each latent factor constituted a random effect and was consequently allowed to vary at Level 2. Thus, variance of the latent factors represented between-person differences in intercept and slopes.

We applied this same two-level latent growth approach to model diurnal cortisol secretion. At Level 1, we specified three latent factors: (1) an intercept that reflects cortisol levels at awakening, (2) a latent response slope capturing the rise following awakening (cortisol awakening response), and (3) a latent slope representing the decline in cortisol from morning to evening (diurnal slope). The model estimated the cortisol awakening response prior to a specified turning point and an independent diurnal slope following the turning point, which was found to be optimal 32 minutes after awakening (Malanchini et al., 2019). Level 1 additionally included a quadratic term (time since turning point squared) to account for non-linearity in the diurnal slope (Miller et al., 2016) and days of sampling (first day, second day) as dummy coded covariates to account for day-to-day variation. See Malanchini et al. (2019) for more information on phenotypic cortisol models.

Lastly, the model of cortisol secretion in response to stress was combined with the diurnal secretion model. At Level 1, we specified five latent factors: (1) a shared intercept that reflected awakening cortisol levels, (2) cortisol awakening response, (3) diurnal slope, (4) response to stress, and (5) recovery following stress.

#### Twin model specification

Behavior genetic models were fit to the data to determine variance attributable to additive genetic influences (*A*), shared environmental influences (*C*), and non-shared environmental influences unique to each twin (*E*; also includes error variance). The *ACE* factors were standardized. Given that monozygotic and dizygotic pairs share approximately 100% and 50% of their segregating genes, respectively, *A* factors were fixed to correlate at 1 for monozygotic pairs and 0.5 for both opposite-sex and same-sex dizygotic pairs. Correlations between *C* factors were fixed to 1 in all twin pairs, given that twins were raised together. Multivariate Cholesky decompositions were conducted in order to examine the extent to which genetic and environmental variance overlapped between measures (see Analytic approach section). These were converted to total genetic correlations (total *r*_A_; Loehlin, 1996). Twin models were run as multigroup models with three groups: monozygotic pairs, dizygotic same-sex pairs, and dizygotic opposite-sex pairs. All models included age (standardized), sex (effect coded as female = −0.5 and male = 0.5), and age-by-sex interaction effects predicting latent cortisol indices.

All models were implemented in *Mplus* 8.2 and were fit with full information maximum likelihood estimation (Muthén & Muthén, 2017). To account for nesting of multiple waves of data within individuals, and multiple twin pairs within families, a sandwich correction was applied to the standard errors in all analyses.

## Results

### Is there genetic variation in cortisol reactions to stress?

We observed significant genetic effects on variation in the pre-stressor cortisol intercept (i.e., baseline cortisol levels prior to acute stress), innovative variation in response to acute in-laboratory stress (i.e., unique of pre-stressor intercept), but no innovative variation in recovery following acute stress (unique of the pre-stressor intercept and stress response; see Table 1 for *ACE* variance estimates without Cholesky decompositions and Table S2 for parameter estimates).

**Table 1.** Combined model of the cortisol reactions to stress and diurnal secretion

| Measure |  | Estimate | S.E. | p | Measure |  | Estimate | S.E. | p |
| --- | --- | --- | --- | --- | --- | --- | --- | --- | --- |
| *Level 1: Within-Person* |  |  |  |  |  |  |  |  |  |
| Day 2 ⇒ Cortisol Values | | 0.095 | 0.025 | <0.001 | | | | | |
| Day 3 ⇒ Cortisol Values | | 0.057 | 0.035 | 0.102 | | | | | |
| Day 4 ⇒ Cortisol Values | | 0.094 | 0.035 | 0.007 | | | | | |
| Day 5 ⇒ Cortisol Values | | 0.119 | 0.046 | 0.010 | | | | | |
| Quadratic Term ⇒ Cortisol Values | | 0.013 | 0.001 | <0.001 | | | | | |
| Sample-Specific Disturbances | A | 0.288 | 0.022 | <0.001 |  |  |  |  |  |
|  | C | 0.000 | 0.000 | 0.747 |  |  |  |  |  |
|  | E | 0.449 | 0.020 | <0.001 |  |  |  |  |  |
| *Level 2: Between-Person Diurnal Secretion* |  |  |  |  |  |  |  |  |  |
| *Cholesky Variance Decomposition* |  |  |  |  | *Total ACE Variance* |  |  |  |  |
| Waking Intercept | A | 0.292 | 0.097 | 0.003 | Waking Intercept | A | 0.292 | 0.097 | 0.003 |
|  | C | 0.480 | 0.071 | <0.001 |  | C | 0.480 | 0.071 | <0.001 |
|  | E | 0.281 | 0.041 | <0.001 |  | E | 0.281 | 0.041 | <0.001 |
| Waking Intercept ⇒ Awakening Response | A | -0.333 | 0.273 | 0.224 | | | | | |
|  | C | -0.279 | 0.180 | 0.122 |  |  |  |  |  |
|  | E | -0.374 | 0.119 | 0.002 |  |  |  |  |  |
| Waking Intercept ⇒ Diurnal Slope | A | -0.007 | 0.013 | 0.586 | | | | | |
|  | C | 0.002 | 0.009 | 0.853 |  |  |  |  |  |
|  | E | -0.002 | 0.005 | 0.777 |  |  |  |  |  |
| Awakening Response Unique of Waking Intercept | A | 0.485 | 0.159 | 0.002 | Awakening Response | A | 0.588 | 0.216 | 0.007 |
|  | C | 0.384 | 0.180 | 0.033 |  | C | 0.474 | 0.222 | 0.033 |
|  | E | 0.389 | 0.071 | <0.001 |  | E | 0.540 | 0.101 | <0.001 |
| Awakening Response Unique of Waking Intercept ⇒ Diurnal Slope | A | 0.001 | 0.012 | 0.919 | | | | | |
|  | C | -0.009 | 0.016 | 0.566 |  |  |  |  |  |
|  | E | -0.002 | 0.006 | 0.676 |  |  |  |  |  |
| Diurnal Slope Unique of Waking Intercept and Awakening Response | A | 0.029 | 0.008 | 0.001 | Diurnal Slope | A | 0.029 | 0.008 | <0.001 |
|  | C | 0.039 | 0.007 | <0.001 |  | C | 0.040 | 0.007 | <0.001 |
|  | E | 0.023 | 0.004 | <0.001 |  | E | 0.023 | 0.004 | <0.001 |
| *Level 2: Between-Person Stress Reaction* |  |  |  |  |  |  |  |  |  |
| *Cholesky Variance Decomposition* |  |  |  |  | *Total ACE Variance* |  |  |  |  |
| Waking Intercept ⇒ Stress Response | A | 0.176 | 0.470 | 0.708 | | | | | |
|  | C | -0.298 | 0.217 | 0.170 |  |  |  |  |  |
|  | E | 0.253 | 0.242 | 0.295 |  |  |  |  |  |
| Waking Intercept ⇒ Stress Recovery | A | 0.042 | 0.151 | 0.781 | | | | | |
|  | C | -0.173 | 0.107 | 0.105 |  |  |  |  |  |
|  | E | 0.046 | 0.078 | 0.558 |  |  |  |  |  |
| Awakening Response Unique of Waking Intercept ⇒ Stress Response | A | 0.074 | 0.432 | 0.865 | | | | | |
|  | C | 0.122 | 0.287 | 0.671 |  |  |  |  |  |
|  | E | -0.040 | 0.241 | 0.869 |  |  |  |  |  |
| Awakening Response Unique of Waking Intercept ⇒ Stress Recovery | A | -0.088 | 0.114 | 0.441 | | | | | |
|  | C | -0.064 | 0.178 | 0.718 |  |  |  |  |  |
|  | E | -0.032 | 0.066 | 0.632 |  |  |  |  |  |
| Diurnal Slope Unique of Waking Intercept and Awakening Response ⇒ Stress | A | -0.655 | 0.393 | 0.095 | | | | | |
|  | C | 0.103 | 0.214 | 0.630 |  |  |  |  |  |
|  | E | -0.002 | 0.200 | 0.992 |  |  |  |  |  |
| Diurnal Slope Unique of Waking Intercept and Awakening Response ⇒ Stress Recovery | A | 0.256 | 0.112 | 0.023 | | | | | |
|  | C | 0.015 | 0.102 | 0.885 |  |  |  |  |  |
|  | E | -0.062 | 0.077 | 0.415 |  |  |  |  |  |
| Stress Response Unique of Waking Intercept, Awakening Response, and Diurnal Slope | A | 1.119 | 0.340 | 0.001 | Stress Response | A | 1.311 | 0.193 | <0.001 |
|  | C | -0.135 | 0.232 | 0.561 |  | C | 0.362 | 0.234 | 0.122 |
|  | E | 1.159 | 0.163 | <0.001 |  | E | 1.188 | 0.182 | <0.001 |
| Stress Response Unique of Waking Intercept, Awakening Response, and Diurnal Slope ⇒ Stress Recovery | A | -0.184 | 0.084 | 0.028 | | | | | |
|  | C | 0.18 | 0.192 | 0.347 |  |  |  |  |  |
|  | E | -0.166 | 0.056 | 0.003 |  |  |  |  |  |
| Stress Recovery Unique of Waking Intercept, Awakening Response, Diurnal Slope, and Stress Response | A | 0.000 | 0.000 | 0.172 | Stress Recovery | A | 0.330 | 0.099 | 0.001 |
|  | C | 0.000 | 0.000 | 0.721 |  | C | 0.258 | 0.120 | 0.031 |
|  | E | 0.000 | 0.000 | 0.555 |  | E | 0.187 | 0.058 | 0.001 |

| *Covariates* |  |  |  |  |
| --- | --- | --- | --- | --- |
| Age ⇒ Waking Intercept | | -0.039 | 0.038 | 0.300 |
| Age ⇒ Awakening Response | | 0.040 | 0.055 | 0.468 |
| Age ⇒ Diurnal Slope | | 0.000 | 0.003 | 0.974 |
| Age ⇒ Stress Response | | 0.239 | 0.102 | 0.019 |
| Age ⇒ Stress Recovery | | -0.060 | 0.045 | 0.179 |
| Sex ⇒ Waking Intercept | | -0.083 | 0.045 | 0.062 |
| Sex ⇒ Awakening Response | | 0.056 | 0.096 | 0.559 |
| Sex ⇒ Diurnal Slope | | -0.001 | 0.006 | 0.874 |
| Sex ⇒ Stress Response | | 0.210 | 0.208 | 0.313 |
| Sex ⇒ Stress Recovery | | 0.115 | 0.085 | 0.179 |
| Age x Sex ⇒ Waking Intercept | | -0.062 | 0.044 | 0.161 |
| Age x Sex ⇒ Awakening Response | | -0.013 | 0.092 | 0.890 |
| Age x Sex ⇒ Diurnal Slope | | 0.005 | 0.006 | 0.386 |
| Age x Sex ⇒ Stress Response | | 0.048 | 0.196 | 0.807 |
| Age x Sex ⇒ Stress Recovery | | 0.070 | 0.076 | 0.356 |
| Stress time ⇒ Awake Intercept | | 0.039 | 0.020 | 0.057 |
| Stress time ⇒ Stress Response | | 0.971 | 0.107 | <0.001 |
| Stress time ⇒ Stress Recovery | | -0.076 | 0.043 | 0.079 |
| *Conditional Means* |  |  |  |  |
| Waking Intercept |  | 1.575 | 0.047 | <0.001 |
| Awakening Response |  | 0.705 | 0.061 | <0.001 |
| Diurnal Slope |  | -0.386 | 0.019 | <0.001 |
| Stress Response |  | 0.412 | 0.118 | <0.001 |
| Stress Recovery |  | -0.591 | 0.051 | <0.001 |

**Model Fit Indices:** -2Log Likelihood = -15412.505, AIC = 30995.009
All estimates are unstandardized. Total ACE Variance = total A, C, or E variance in each outcome accounted for by all respective components of variance. Stress time = time between waking and start of TSST-C. ⇒ Cholesky path. Age was standardized, sex was effect coded. Units are in raw concentrations residualized for batch year and log-transformed.

These three components of genetic and phenotypic variation are illustrated in **Figure 1**. The blue, black, and red lines represent expected phenotypic trajectories for individuals who were, respectively,1 *SD* above the mean, at the mean, and 1 *SD* below the mean on genetic dispositions for the following variance components: pre-stressor cortisol levels (A), acute stress responses (B), and stress recovery (C). The trajectories represent the expected mean cortisol trajectories, stratified by level of genetic disposition on each variance component, allowing for effects to magnify or diminish over time, as indicated by the dependencies among the components. These expected means by genotype are superimposed upon the full +/- 1 *SD* phenotypic range of variation in the respective variance components.

**Figure 1.**
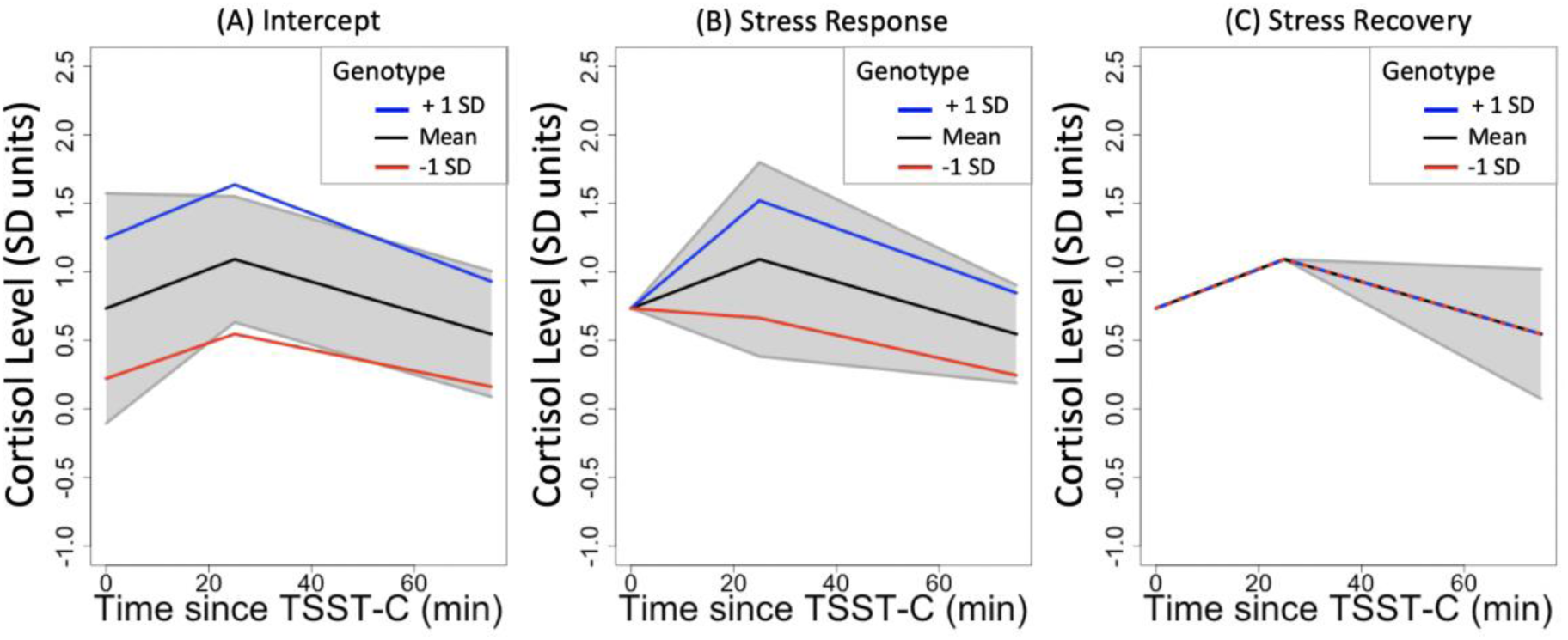
Patterns of individual differences in cortisol reactions to stress accounted for by genetic variability. *Note.* The blue, black, and red lines represent expected trajectories for individuals who were higher (1 *SD* above the mean), average, and lower (1 *SD* above the mean), respectively, on genetic dispositions for pre-stressor cortisol levels and its downstream genetic effects on stress response and recovery (A), stress responses unique of the intercept and its downstream genetic effects on stress recovery (B), and stress recovery unique of the intercept and stress response (C). These expected means by genotype are superimposed upon the full +/- 1 *SD* phenotypic range of variation indicated by the gray shading in the respective variance components. Raw cortisol levels were residualized for assay batch and log-transformed.

Panel A of Figure 1 illustrates genotype differences in baseline cortisol levels prior to acute stress exposure and how such differences progress over the course of the stressor protocol. There were sizable differences in baseline cortisol levels that were strongly heritable and persisted across the hour-long laboratory assessment. Baseline genetic differences in cortisol levels were not related to genetic differences in stress response, and therefore no slope differences were observed between genotypes over the first ∼20 minutes. In contrast, baseline genetic differences were negatively associated with cortisol recovery, as indicated by the subtle narrowing of cortisol differences associated with genotypes over the last ∼40 minutes of the protocol.

Panel B of Figure 1 illustrates the sizable magnitude of genetic effects on the cortisol response to acute stress that were independent of genetic effects on baseline levels. Individuals with similar genotypes for elevated pre-stress baseline levels subsequently diverged phenotypically in response to a stressful environment, as largely independent genotypes for stress response magnitude were evoked.

This is further depicted in **Figure 2**, where the blue, black, and red lines represent expected phenotypic trajectories for individuals who were higher (1 *SD* above the mean), average, and lower (1 *SD* below the mean) respectively on genetic dispositions for the pre-stress intercept. The trajectories diverged into solid and dashed lines, because genotypes for higher (1 *SD* above the mean) and lower (1 *SD* above the mean) stress responses were innovative. Hence, the shaded areas estimate reaction ranges of different genotypes that were substantial enough to reorder individual differences during the course of stressor exposure.

**Figure 2.**
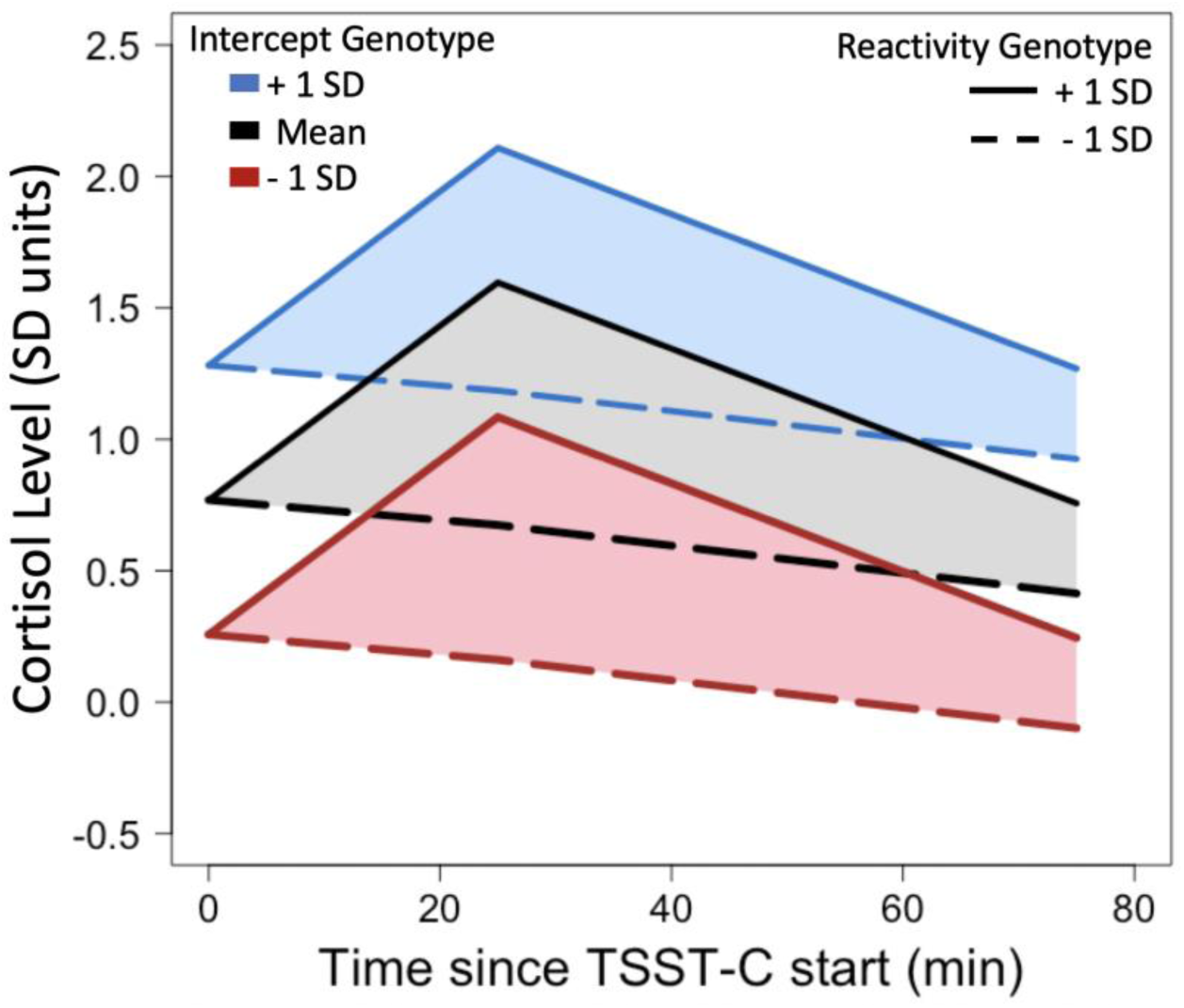
Reaction ranges in cortisol responses to stress reorder individual differences in cortisol output. *Note.* The first cortisol value at 0 minutes is the pre-stress intercept. The blue, black, and red lines represent expected trajectories for individuals who were higher (1 SD above the mean), average, and lower (1 SD below the mean) on genetic dispositions for the pre-stress intercept, respectively. The trajectories diverge into solid and dashed lines as genotypes for higher (1 SD above the mean) and lower (1 SD above the mean) stress responses were innovative. The shaded areas depict the range of reactivity of different genotypes. Raw cortisol levels were residualized for assay batch and log-transformed.

### Is there genetic variation in cortisol change over the course of the day?

We observed significant genetic effects on variation in the cortisol intercept at awakening, innovative variation in cortisol awakening response (unique of intercept), and innovative variation in diurnal slope (unique of intercept and awakening response; see Table 1 for *ACE* variance estimates without Cholesky decompositions and Table S3 for parameter estimates).

These three components of genetic and phenotypic variation are illustrated in **Figure 3**. The blue, black, and red lines represent expected phenotypic trajectories for individuals who were, respectively, 1 *SD* above the mean, at the mean, and 1 *SD* below the mean on genetic dispositions for the following variance components: cortisol intercept at awakening (A), cortisol awakening response (B), and diurnal slope (C). The trajectories represent the expected mean cortisol trajectories stratified by level of genetic disposition on each component of variance (intercept, cortisol awakening response, and diurnal slope), allowing for effects to magnify or diminish over time, as indicated by the dependencies among the components. These expected means by genotype are superimposed upon the full +/- 1 SD phenotypic range of variation in the respective variance components.

**Figure 3.**
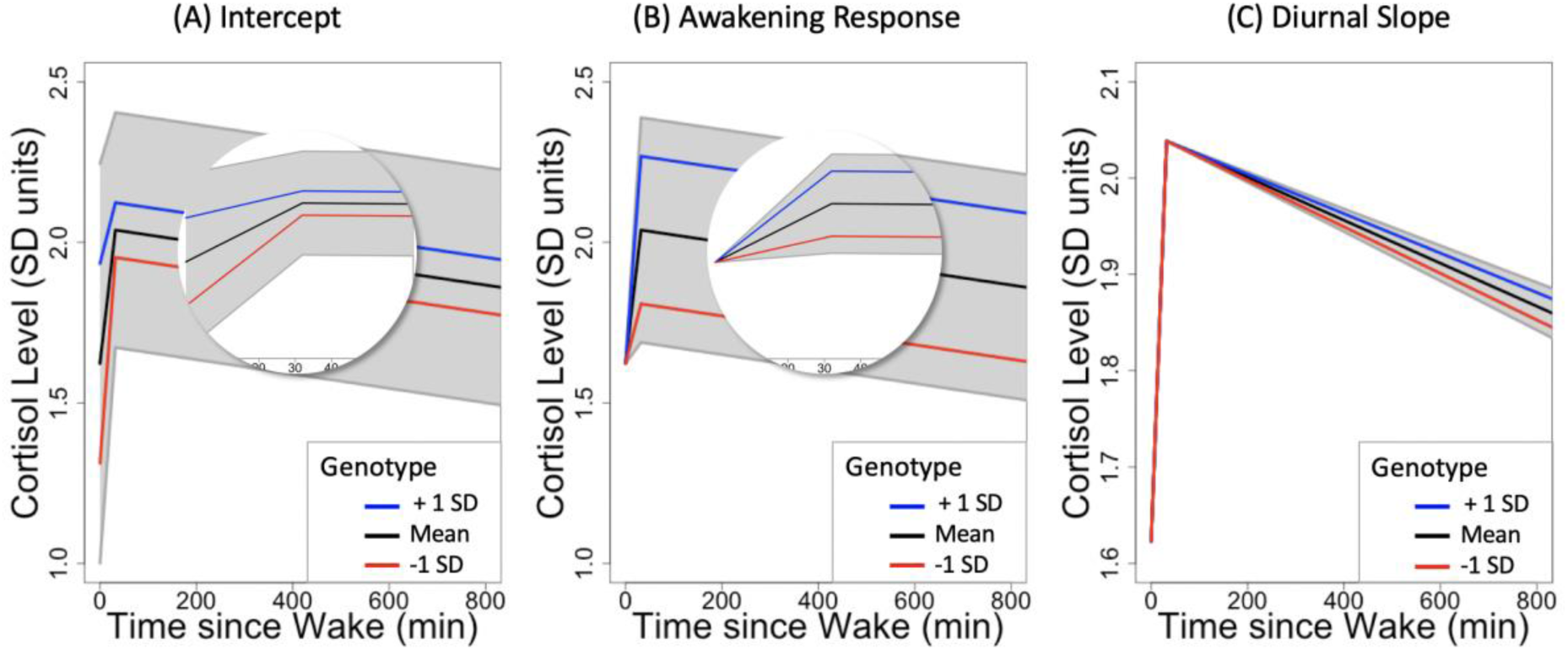
Patterns of individual differences in diurnal cortisol secretion accounted for by genetic variability. *Note.* The blue, black, and red lines represent expected trajectories for individuals who were higher (1 SD above the mean), average, and lower (1 SD above the mean) respectively on genetic dispositions for awakening cortisol intercept and its downstream genetic effects on awakening responses and diurnal slopes (A), awakening response unique of the intercept and its downstream genetic effects on the diurnal slope (B), and the diurnal slope unique of the intercept and awakening response (C). These expected means by genotype are superimposed upon the full +/- 1 SD phenotypic range of variation indicated by the gray shading in the respective variance components. See full text for further interpretation. Circles are zoomed in on the first hour after awakening. Y-axis scaling for C differs to aid visibility of diurnal slope effects. Raw cortisol levels were residualized for assay batch and log-transformed.

Panel A of Figure 3 illustrates genotype differences in baseline cortisol levels at awakening and how such differences progress over the course of the day. There were sizable differences in baseline cortisol levels at awakening that were strongly heritable. Baseline genetic differences at awakening were negatively associated with the awakening response, as indicated by the cortisol differences associated with genotypes over the first ∼60 minutes of the day. Individuals with genetic dispositions for higher awakening levels subsequently showed *lower* awakening responses (blue line) than individuals with genetic dispositions for lower awakening levels (red line). Therefore, following the cortisol awaking response, genotypes related to substantially higher levels at awakening were associated with only slightly higher subsequent cortisol levels throughout the day.

Panel B of Figure 3 illustrates the sizable magnitude of genetic effects on the cortisol response to awakening that were independent of genetic effects on baseline awakening levels. Individuals with similar genotypes for elevated awakening levels subsequently diverged phenotypically in response to awakening, as largely independent genotypes for awakening magnitude were evoked. Individuals with genetic dispositions for higher awakening responses subsequently showed higher cortisol levels (blue line) than individuals with genetic dispositions for lower awakening responses (red line). Thus, genotypes for higher cortisol levels at awakening cannot be used to infer subsequent cortisol levels across the day, because innovative genotypes are evoked in response to awakening.

Panel C of Figure 3 illustrates the sizable magnitude of genetic effects on the diurnal slope that were independent of genetic effects on baseline awakening levels and awakening responses. Genetic differences in baseline cortisol levels and awakening responses were not related to genetic differences in the diurnal slope, and therefore no slope differences were observed between genotypes after ∼30 minutes in panel A and B.

### Are cortisol reactions to stress and the day regulated by the same genetic variation?

The total genetic correlation between the diurnal slope and the cortisol response to stress (total *r*_A_ = −0.516, *SE* = 0.319, *p* = 0.106) or the recovery following stress (total *r*_A_ = 0.280, *SE* = 0.284, *p* = 0.324) were modest to moderate and not reliably different from zero. The magnitude of these and other genetic correlations is depicted in **Figure 4**.

**Figure 4.**
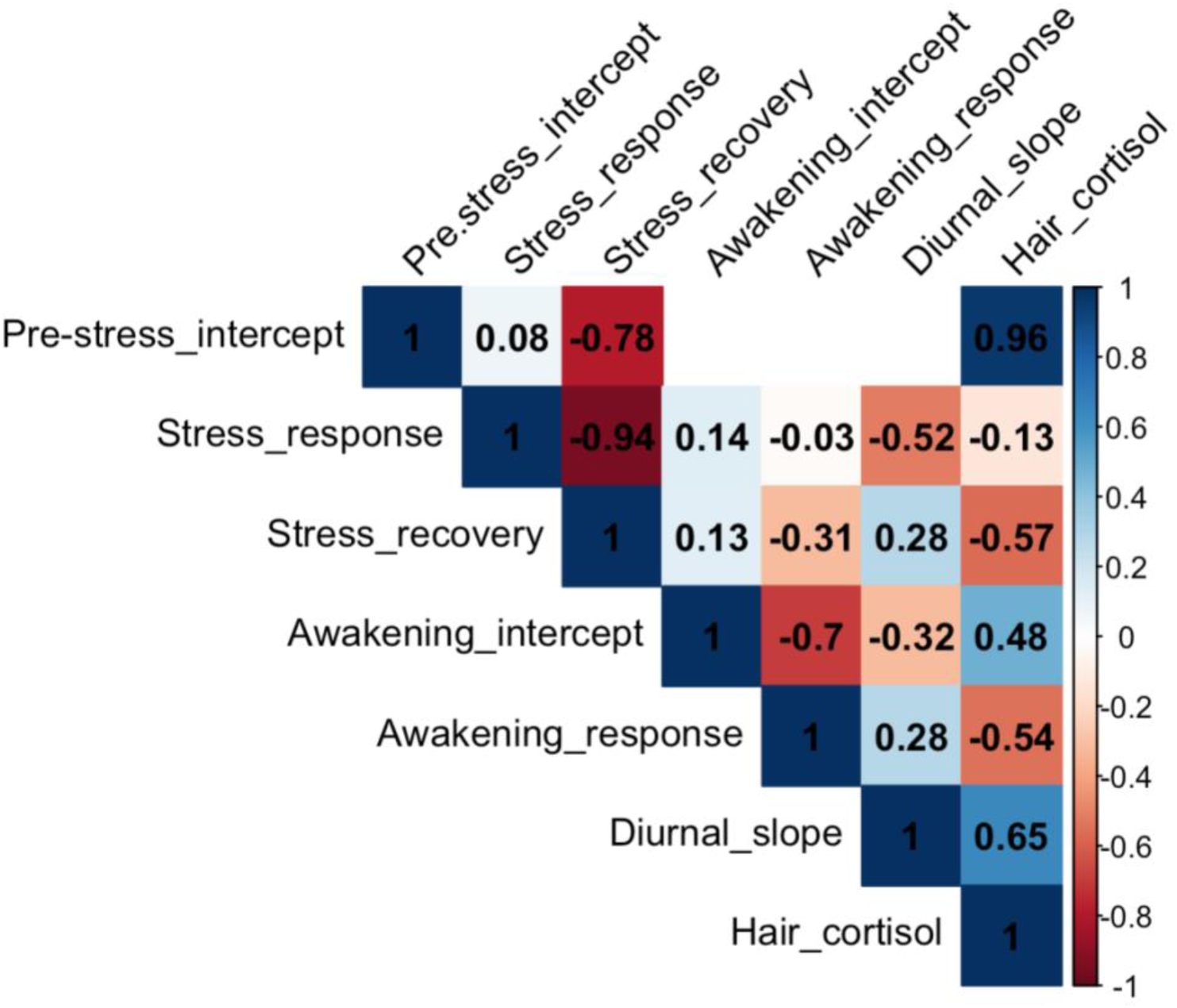
Total genetic correlations between cortisol reactions to stress, diurnal secretion, and hair cortisol. *Note.* Genetic correlations were computed on the basis of 5 separate models: (1) acute stress reaction only, (2) diurnal secretion only, (3) combined model of acute stress reaction and diurnal secretion, (4) acute stress reaction with hair cortisol, and (5) diurnal secretion with hair cortisol. There were no correlations of the pre-stress intercept with diurnal secretion (white cubes), because the combined model includes only one intercept.

The genetic correlation between the cortisol awakening response with cortisol response to stress (total *r*_A_ = −0.030, *SE* = 0.277, *p* = 0.912) or recovery following stress (total *r*_A_ = −0.313, *SE* = 0.245 *p* = 0.202) was negligible to modest. Critically, a significant genetic effect on the cortisol response to stress unique of awakening intercept, awakening response, and diurnal slope was still found (see **Table 1**). Therefore, the genetic variation involved in reactions to stress and the day were largely uncorrelated.

### Are cortisol reactions to stress and the day regulated by the same genetic variance as hair cortisol levels?

The genetic correlations of hair cortisol levels with the cortisol pre-stress intercept was high (total *r*_A_ = 0.965, *SE* = 0.189, *p* < 0.001, see Figure 4). In contrast, genetic correlations of hair cortisol with the response to stress (total *r*_A_ = −0.133, *SE* = 0.760, *p* = 0.861) or recovery following stress (total *r*_A_ = −0.566, *SE* = 0.795, *p* = 0.477) were negligible to moderate and not reliably different from zero. Hence, a changing stressful environment evoked innovative genetic variation and did not just amplify individual differences in hair levels. Notably, genetic effects on intercept, stress response, and recovery unique of hair cortisol and each other were no longer reliably different from zero. See Supplemental Table S4 for full parameter estimates.

Next, genetic correlations of hair cortisol levels with the cortisol intercept at awakening (total *r*_A_ = 0.485, *SE* = 0.398, *p* = 0.223), cortisol awakening response (total *r*_A_ = −0.544, *SE* = 0.478, *p* = 0.255), or diurnal slope (total *r*_A_ = 0.647, *SE* = 0.388, *p* = 0.096) were modest to moderate and not reliably different from zero. Thus, cortisol secretion across the day evoked innovative genetic variation in salivary cortisol secretion and did not just amplify individual differences in hair levels. Notably, a genetic effect of the awakening intercept unique of hair cortisol was still found to be reliably different from zero, but was not true for the awakening response (unique of hair cortisol and awakening intercept) or diurnal slope (unique of hair cortisol, awakening intercept, and awakening response). See Supplemental Table S5 for parameter estimates.

These findings suggest that hair cortisol levels capture some of the genetic variation of stress reactions and diurnal secretion. At the same time, the lack of high genetic correlations that reliably differ from zero indicates that hair cortisol levels do not fully capture the genetic variation involved in dynamic changes.

## Discussion

We present results from the most comprehensive behavioral genetic study of multimodal cortisol response to date. Results indicated that genetic variation was associated with dynamic patterns of cortisol secretion, both in response to a standardized in-laboratory stressor and across the day. Counter to the view that environmental effects either compete with genetic effects or merely serve to magnify standing genetic influences on biopsychosocial phenotypes, cortisol responses to acute stress were regulated by innovative genetic variation that was not apparent prior to stressor onset or in chronic hair cortisol levels. Second, genetic variation in cortisol changes in response to acute stress was largely uncorrelated with genetic variation in cortisol changes across the day (*e.g.*, the genetic correlation of the response to stress and response to awakening was negligible).

Many previous studies of gene × environment interactions on biopsychosocial phenotypes have been limited by comparing different groups of individuals in different environmental contexts, rather than identifying individual differences in within-person change over time, and by the lack of experimental control over environmental change. The current study advances the literature by examining genetic variation in within-person change in cortisol in two contexts: (1) an exogenously-imposed environmental stressor administered in a controlled laboratory condition, and (2) naturally occurring changes throughout the day. In both contexts, mean changes in cortisol secretion were associated with a substantial reordering of individuals, partly on the basis of their genotypes.

The fact that we identified genetic factors relevant to cortisol change that were independent of those relevant to baseline and chronic levels of cortisol variation indicates that it would be inappropriate to describe one genotype or another as coding for higher cortisol output. Rather, the relative ordering of people in their cortisol levels was dependent on the context. This pattern resembles classic findings on genetic reaction ranges in fruit flies (Gupta & Lewontin, 1982) and mice (Cooper & Zubek, 1958), in which relative ordering of organisms on multiple traits substantially changed across environments. Here, we provide an empirical demonstration of the same theoretical process in a behaviorally-relevant human phenotype. Despite some convergence of genetic effects across environments (stress and day), innovative genetic variation was evoked in response to a new stressful environment. Over the course of awakening, the same genetic variation had inverse effects over time. Similarly, across modalities of cortisol secretion, genetic variation of hair cortisol levels was largely independent of genetic variation involved in salivary responses to stress or the day.

These findings contradict the notion that environmental change or intervention merely amplifies pre-existing genetic differences. Rather, these results provide evidence that changing environments can evoke genetic variation that might remain silent in alternative situations or even evoke inverse effects of standing genetic variation. As a result, genetically-associated differences observed in one group in a specific environment do not fully inform the relative ordering of genetically-associated individual differences of that group in a new environment (Gottlieb, 2007), or what differences between groups will be in a new environment (Taylor, 2006).

Cortisol secretion is a model phenotype that is well-suited for the study of gene × environment interactions because of its behavioral relevance, responsiveness to change over short timescales, and heritability. Despite these strengths, our results may not generalize to other psychological domains that develop and change more gradually, such as cognitive ability. Evaluating the generalizability of our findings that show innovative genetic variation in changing environments will require further genetically-informative studies that exogenously manipulate environments and characterize interactively changing reactions to them. For instance, future studies could explore genetic variation in behaviorally or neuroanatomically observed learning curves of new knowledge (*e.g.*, an unfamiliar language) and new skills (*e.g.*, writing with the non-dominant hand, learning a new musical instrument). Such studies could employ twin-based designs, as was applied here, and/or they could capitalize on genome-wide molecular genetic measures (*e.g.*, polygenic scores; Belsky & Harden, 2019).

In conclusion, this study provides empirical evidence that the genome regulates individuals’ reactions to the environment that differ across environments. If environments are constantly changing, it follows that the genetic factors that are relevant to the outcomes under study may be continuously in flux.

## Acknowledgments

We gratefully acknowledge all participants of the Texas Twin Project. This research was supported by NIH grant R01HD083613 and R01HD092548. LR is supported by the German Research Foundation (DFG). KPH and EMTD are Faculty Research Associates of the Population Research Center at the University of Texas at Austin, which is supported by a grant, 5-R24-HD042849, from the Eunice Kennedy Shriver National Institute of Child Health and Human Development (NICHD). KPH and EMTD are also supported by Jacobs Foundation Research Fellowships.

## Author Contributions

KPH, EMTD, MM, & LR developed the study concept and design. Testing and data collection were performed by AG, AS, JM, CY, MP, LE. LR & MM performed the data analysis and interpretation under the supervision of EMTD and KPH. LR drafted the manuscript. All authors provided critical revisions and approved the final version of the manuscript for submission.

## Open Practices Statement

Analyses reported in this article were not formally preregistered. The data have not been made available on a permanent third-party archive; requests for the model scripts can be sent via email to the lead author at.

