## Supplement for "Changing environments reveal innovative genetic variation in children’s cortisol responses"

### Supplemental Method

#### *Cortisol Stress Response*

Participants were instructed to refrain from eating 1 hr before their lab visit. Twin pairs came to the lab together, but each child completed the TSST–C separately. Salivary samples were collected as passive drool extracted into 2 mL plastic vials. Research assistants recorded the exact time at which each sample was collected. All samples were frozen at the same time (maximum of 2.5 hrs from the collection of the first sample) at -80 degrees Fahrenheit prior to being shipped on dry ice to Dr. Clemens Kirschbaum's lab in Germany.

#### *Diurnal Cortisol Secretion*

Participants were asked to refrain from eating, drinking, or brushing their teeth for the 30 min preceding each sample, and they were provided with diaries where they could record their daily activities and experiences regarding the data collection. Participants were instructed to place each vial in their home freezer immediately after sampling. Saliva samples were returned to the lab the day after saliva collection was completed using a provided pre–paid envelope. Samples were frozen -80 degrees Fahrenheit in the lab prior to being shipped on dry ice to Dr. Clemens Kirschbaum's laboratory.

Minor deviations in sampling timing can have dramatic effects on cortisol values, especially in the morning (Stalder et al., 2015). Thus, each sampling vial had to be removed from a bottle equipped with an electronic date- and time-tracking cap (MEMs Track Cap; Aardex, Denver, CO), and participants recorded the date and time of collection on an adhesive label attached to each vial after sampling. MEMs cap times were unavailable for 13.73% of returned samples, primarily due to product failure or because participants had opened the MEMs container to remove samples 1 and 2 simultaneously. For those with MEMs data available, the median deviation of MEMs-recorded time from that reported by participants ranged from 2-4 minutes across all samples and days (mean = 6.50, SD = 9.24). Participant- or parent-reported sampling times were used for the current quality control procedures and following analyses; if those were missing, the MEMs cap times were used.

Individual samples were excluded when they deviated severely from normative diurnal secretory patterns, indicating a failure to provide saliva at the correct sampling times or an abnormal sleep–wake schedule. Sample 1 was excluded when the interval between reported wake and sample 1 exceeded 20 minutes, as this is considered not indicative of waking levels, but of the cortisol awakening response (*n* ranging from 0 samples on day 3 to 8 for day 2). Sample 2 was excluded when the time interval between sample 1 and 2 exceeded 60 minutes (daily *n* ranged from 2 to 7 across the 5 days of data collection). Sample 2 was also excluded when it was lower than concentrations for the evening sample on the same day (*n* excluded day1 = 30, *n* excluded day 2 = 29, *n* excluded day 3 = 23, *n* excluded day 4 = 32, *n* excluded day 5 = 4). Lastly, sample 3 was excluded when its concentrations were higher than the next day's waking concentrations (*n* excluded day1-2 = 35, *n* excluded day 2-3 = 29, *n* excluded day 3-4 = 37)

##### ***Hair Cortisol***

Participants were instructed to refrain from using leave-in hair products, such as hair gel, on the day of the lab visit. Samples were stored in a dry location and shipped to Dr. Clemens Kirschbaum's lab for steroid measurement. Technical details on the extraction procedure are provided elsewhere Gao et al., 2013. Internal consistency estimates for cortisol analyzed using liquid chromatography tandem mass spectrometry (LC-MS/MS) have been reported above 0.96 (Stalder et al., 2012). The lower limit of sensitivity for hair cortisol was 0.1 pg/ml (Gao et al., 2013).

##### ***Zygosity***

Opposite-sex twin pairs were classified as dizygotic. Same-sex twin pair zygosity was assessed using responses to ratings about the twins' physical similarities (e.g., facial appearance). The ratings were completed by parents and two research assistants. Parents additionally rated how often the twins are mistaken for one another. These ratings were entered into a latent class analysis that was used to obtain zygosity classifications. Latent class analysis has been reported to accurately determine DNA-based zygosity >99% of the time (Heath et al., 2003). In the present study, latent class analysis accurately determined zygosity >97% of the time in 713 genotyped individuals.

#### Supplemental Tables

**Table S1.** Descriptive statistics for raw cortisol values.
**Table S2.** Latent growth ACE model of cortisol reactions to stress.
**Table S3.** Latent growth ACE model of diurnal secretion.
**Table S4.** Latent growth ACE model of cortisol stress response combined with hair cortisol.
**Table S5.** Latent growth ACE model of diurnal cortisol secretion combined with hair cortisol.

Table S1. Descriptive statistics for raw cortisol values.

| Sample | *n* | *M* | *SD* |
|---|---|---|---|
| Stress Cortisol 1 | 535 | 5.61 | 47.25 |
| Stress Cortisol 2 | 519 | 6.45 | 39.38 |
| Stress Cortisol 3 | 519 | 4.40 | 17.16 |
| Stress Cortisol 4 | 518 | 3.52 | 16.22 |
| Diurnal Cortisol 1, day 1 | 550 | 7.70 | 17.30 |
| Diurnal Cortisol 2, day 1 | 504 | 13.25 | 83.19 |
| Diurnal Cortisol 3, day 1 | 500 | 0.96 | 5.21 |
| Diurnal Cortisol 1, day 2 | 566 | 7.27 | 11.23 |
| Diurnal Cortisol 2, day 2 | 521 | 9.67 | 7.27 |
| Diurnal Cortisol 3, day 2 | 525 | 0.79 | 2.06 |
| Diurnal Cortisol 1, day 3 | 574 | 6.94 | 7.20 |
| Diurnal Cortisol 2, day 3 | 526 | 9.57 | 8.19 |
| Diurnal Cortisol 3, day 3 | 502 | 0.67 | 1.50 |
| Diurnal Cortisol 1, day 4 | 566 | 6.78 | 5.14 |
| Diurnal Cortisol 2, day 4 | 506 | 8.95 | 8.04 |
| Diurnal Cortisol 3, day 4 | 559 | 1.34 | 4.70 |
| Diurnal Cortisol 1, day 5 | 126 | 7.67 | 9.09 |
| Diurnal Cortisol 2, day 5 | 96 | 30.88 | 217.28 |
| Diurnal Cortisol 3, day 5 | 110 | 1.5 | 4.09 |
| Hair Cortisol | 1338 | 11.09 | 68.61 |

Note: Descriptive statistics for raw cortisol values after exclusions and before cortisol residualization for year of assay and log transformation. Salivary cortisol concentrations are measured in nmol/L and hair cortisol concentrations in pg/ml. Diurnal and hair cortisol descriptive statistics include repeat participants.

Table S2. Latent growth ACE model of cortisol reactions to stress.

| **Measure** | | **Estimate** | **S.E.** | ***p*** |
|---|---|---|---|---|
| *Level 1: Within-Person* | | | | |
| Sample-Specific Disturbances | A | 0.025 | 0.315 | 0.937 |
| | C | 0.062 | 0.090 | 0.489 |
| | E | 0.201 | 0.022 | <0.001 |
| *Level 2: Between-Person Variance Decomposition* | | | | |
| Pre-Stress Intercept | A | 0.512 | 0.093 | <0.001 |
| | C | 0.188 | 0.156 | 0.228 |
| | E | 0.637 | 0.072 | <0.001 |
| Pre-Stress Intercept ⇒ Stress Response | A | 0.080 | 0.379 | 0.833 |
| | C | -0.801 | 0.205 | <0.001 |
| | E | -1.028 | 0.170 | <0.001 |
| Pre-Stress Intercept ⇒ Stress Recovery | A | -0.194 | 0.129 | 0.133 |
| | C | 0.390 | 0.094 | <0.001 |
| | E | 0.040 | 0.061 | 0.513 |
| Stress Response Unique of Pre-Stress Intercept | A | 1.028 | 0.314 | 0.001 |
| | C | 0.000 | 0.009 | 0.996 |
| | E | 1.351 | 0.106 | <0.001 |
| Stress Response Unique of Pre-Stress Intercept ⇒ Stress Recovery | A | -0.154 | 0.113 | 0.173 |
| | C | 0.000 | 0.005 | 0.996 |
| | E | -0.412 | 0.051 | <0.001 |
| Stress Recovery Unique of Pre-Stress Intercept and Stress Response | A | 0.000 | 0.000 | 0.349 |
| | C | 0.000 | 0.000 | 0.076 |
| | E | 0.567 | 0.040 | <0.001 |
| *Covariates* | | | | |
| Age ⇒ Pre-TSST Intercept | | 0.153 | 0.039 | <0.001 |
| Age ⇒ Stress Response | | -0.030 | 0.096 | 0.752 |
| Age ⇒ Stress Recovery | | -0.102 | 0.040 | 0.010 |
| Sex ⇒ Pre-TSST Intercept | | 0.041 | 0.079 | 0.603 |
| Sex ⇒ Stress Response | | -0.574 | 0.208 | 0.006 |
| Sex ⇒ Stress Recovery | | 0.132 | 0.081 | 0.102 |
| Age x Sex ⇒ Pre-TSST Intercept | | -0.054 | 0.062 | 0.381 |
| Age x Sex ⇒ Stress Response | | -0.177 | 0.184 | 0.336 |
| Age x Sex ⇒ Stress Recovery | | 0.111 | 0.072 | 0.121 |
| Pre-stress time ⇒ Pre-TSST Intercept | | -0.079 | 0.040 | 0.048 |
| Pre-stress time ⇒ Stress Response | | 0.259 | 0.101 | 0.010 |
| Pre-stress time ⇒ Stress Recovery | | -0.072 | 0.041 | 0.081 |
| *Intercepts* | | | | |
| Pre-Stress Intercept | | 0.769 | 0.048 | <0.001 |
| Stress Response | | 0.878 | 0.113 | <0.001 |
| Stress Recovery | | -0.660 | 0.044 | <0.001 |
| **Model Fit Indices:** -2Log Likelihood = -3609.866, AIC = 7309.732 | | | | |

All estimates are unstandardized. Units are in raw concentrations residualized for batch year and log-transformed. ⇒ Cholesky path; Pre-stress time = time between waking and start of TSST-C. Age and pre-stress time were standardized; sex was effect coded.

Table S3. Latent growth ACE model of diurnal secretion.

| **Measures** | | **Estimate** | **S.E.** | ***p*** |
|---|---|---|---|---|
| *Level 1: Within-Person* | | | | |
| Day 2 ⇒ Cortisol Values | | 0.047 | 0.026 | 0.068 |
| Day 3 ⇒ Cortisol Values | | 0.010 | 0.037 | 0.779 |
| Day 4 ⇒ Cortisol Values | | 0.046 | 0.040 | 0.247 |
| Day 5 ⇒ Cortisol Values | | 0.063 | 0.045 | 0.166 |
| Quadratic Term ⇒ Cortisol Values | | 0.009 | 0.002 | <0.001 |
| Sample-Specific Disturbances | A | 0.271 | 0.029 | <0.001 |
| | C | 0.000 | 0.000 | 0.425 |
| | E | 0.471 | 0.023 | <0.001 |
| *Level 2: Between-Person Variance Decomposition* | | | | |
| Waking Intercept | A | 0.311 | 0.103 | 0.002 |
| | C | 0.467 | 0.077 | <0.001 |
| | E | 0.269 | 0.042 | <0.001 |
| Waking Intercept ⇒ Awakening Response | A | -0.423 | 0.281 | 0.133 |
| | C | -0.137 | 0.222 | 0.537 |
| | E | -0.384 | 0.122 | 0.002 |
| Waking Intercept ⇒ Diurnal Slope | A | -0.009 | 0.013 | 0.461 |
| | C | 0.000 | 0.008 | 0.995 |
| | E | 0.001 | 0.006 | 0.903 |
| Awakening Response Unique of Waking Intercept | A | 0.431 | 0.189 | 0.022 |
| | C | 0.336 | 0.288 | 0.242 |
| | E | 0.364 | 0.081 | <0.001 |
| Awakening Response Unique of Waking Intercept ⇒ Diurnal Slope | A | 0.002 | 0.013 | 0.849 |
| | C | -0.002 | 0.019 | 0.923 |
| | E | 0.004 | 0.005 | 0.505 |
| Diurnal Slope Unique of Waking Intercept and Awakening Response | A | 0.028 | 0.010 | 0.008 |
| | C | 0.035 | 0.007 | <0.001 |
| | E | 0.019 | 0.006 | 0.001 |
| *Covariates* | | | | |
| Age ⇒ Waking Intercept | | -0.046 | 0.035 | 0.197 |
| Age ⇒ Awakening Response | | -0.003 | 0.055 | 0.957 |
| Age ⇒ Diurnal Slope | | 0.001 | 0.003 | 0.747 |
| Sex ⇒ Waking Intercept | | -0.088 | 0.041 | 0.031 |
| Sex ⇒ Awakening Response | | -0.165 | 0.086 | 0.056 |
| Sex ⇒ Diurnal Slope | | 0.002 | 0.005 | 0.645 |
| Age x Sex ⇒ Waking Intercept | | 0.001 | 0.003 | 0.747 |
| Age x Sex ⇒ Awakening Response | | -0.083 | 0.067 | 0.214 |
| Age x Sex ⇒ Diurnal Slope | | -0.003 | 0.006 | 0.577 |
| *Conditional Means* | | | | |
| Waking Intercept | | 1.623 | 0.047 | <0.001 |
| Awakening Response | | 0.779 | 0.056 | <0.001 |
| Diurnal Slope | | -0.343 | 0.021 | <0.001 |
| **Model Fit Indices:** -2Log Likelihood = -10999.257, AIC = 22084.515 | | | | |

All estimates are unstandardized. Units are in raw concentrations residualized for batch year and log-transformed; ⇒ Cholesky path. Age was standardized; sex was effect coded.

Table S4. Latent growth ACE model of cortisol stress response combined with hair cortisol.

| **Measure** | | **Estimate** | **S.E.** | ***p*** |
|---|---|---|---|---|
| *Level 1: Within-Person* | | | | |
| Sample-Specific Disturbances | A | 0.024 | 0.326 | 0.942 |
| | C | 0.062 | 0.089 | 0.481 |
| | E | 0.201 | 0.022 | <0.001 |
| *Level 2: Between-Person Variance Decomposition* | | | | |
| Hair Cortisol | A | 0.203 | 0.179 | 0.257 |
| | C | 0.731 | 0.100 | <0.001 |
| | E | 0.770 | 0.052 | <0.001 |
| Hair Cortisol ⇒ Pre-Stress Intercept | A | 0.447 | 0.261 | 0.087 |
| | C | 0.130 | 0.15 | 0.387 |
| | E | 0.048 | 0.071 | 0.493 |
| Pre-Stress Intercept Unique of Hair Cortisol | A | -0.122 | 0.315 | 0.698 |
| | C | 0.220 | 0.307 | 0.474 |
| | E | 0.645 | 0.087 | <0.001 |
| Hair Cortisol ⇒ Stress Response | A | -0.143 | 0.824 | 0.862 |
| | C | -0.272 | 0.215 | 0.206 |
| | E | -0.011 | 0.122 | 0.927 |
| Pre-Stress Intercept Unique of Hair Cortisol ⇒ Stress Response | A | -1.067 | 0.27 | <0.001 |
| | C | -0.592 | 0.702 | 0.399 |
| | E | -1.023 | 0.167 | <0.001 |
| Stress Response Unique of Pre-Stress Intercept and Hair Cortisol | A | 0.000 | 0.000 | 0.964 |
| | C | -0.394 | 0.998 | 0.693 |
| | E | 1.343 | 0.105 | <0.001 |
| Hair Cortisol ⇒ Stress Recovery | A | -0.154 | 0.232 | 0.508 |
| | C | 0.139 | 0.081 | 0.085 |
| | E | 0.014 | 0.041 | 0.728 |
| Pre-Stress Intercept Unique of Hair Cortisol ⇒ Stress Recovery | A | 0.224 | 0.16 | 0.160 |
| | C | 0.245 | 0.576 | 0.670 |
| | E | 0.034 | 0.069 | 0.623 |
| Stress Response Unique of Pre-Stress Intercept and Hair Cortisol ⇒ Stress Recovery | A | 0.000 | 0.000 | 0.908 |
| | C | 0.249 | 0.618 | 0.687 |
| | E | -0.411 | 0.051 | <0.001 |
| Stress Recovery Unique of Pre-Stress Intercept, Stress Response, and Hair Cortisol | A | 0.000 | 0.000 | 0.411 |
| | C | 0.000 | 0.000 | 0.076 |
| | E | 0.567 | 0.041 | <0.001 |
| *Covariates* | | | | |
| Age ⇒ Hair | | -0.059 | 0.051 | 0.242 |
| Age ⇒ Pre-TSST Intercept | | 0.147 | 0.039 | <0.001 |
| Age ⇒ Stress Response | | -0.025 | 0.095 | 0.789 |
| Age ⇒ Stress Recovery | | -0.104 | 0.040 | 0.009 |
| Sex ⇒ Hair | | 0.332 | 0.092 | <0.001 |
| Sex ⇒ Pre-TSST Intercept | | 0.052 | 0.075 | 0.486 |
| Sex ⇒ Stress Response | | -0.595 | 0.207 | 0.004 |
| Sex ⇒ Stress Recovery | | 0.141 | 0.081 | 0.081 |

| | | | |
|---|---|---|---|
| Age x Sex ⇒ Hair | 0.02 | 0.098 | 0.839 |
| Age x Sex ⇒ Pre-TSST Intercept | -0.069 | 0.062 | 0.266 |
| Age x Sex ⇒ Stress Response | -0.148 | 0.18 | 0.411 |
| Age x Sex ⇒ Stress Recovery | 0.100 | 0.071 | 0.157 |
| Pre-stress time ⇒ Pre-TSST Intercept | -0.085 | 0.037 | 0.022 |
| Pre-stress time ⇒ Stress Response | 0.264 | 0.101 | 0.009 |
| Pre-stress time ⇒ Stress Recovery | -0.073 | 0.041 | 0.075 |
| *Conditional Means* | | | |
| Pre- Stress Intercept | 0.766 | 0.047 | <0.001 |
| Stress Response | 0.883 | 0.112 | <0.001 |
| Stress Recovery | -0.662 | 0.044 | <0.001 |
| Hair | 0.095 | 0.053 | 0.075 |
| **Model Fit Indices:** -2Log Likelihood = -4453.334, AIC = 9028.668 | | | |

All estimates are unstandardized. Units are in raw concentrations residualized for batch year and log-transformed. ⇒ Cholesky path; Pre-stress time = time between waking to start of TSST–C. Age and stress time were standardized; sex was effect coded.

Table S5. Latent growth ACE model of diurnal cortisol secretion combined with hair cortisol.

| **Measures** | | **Estimate** | **S.E.** | ***p*** |
|---|---|---|---|---|
| *Level 1: Within-Person* | | | | |
| Day 2 ⇒ Cortisol Values | | 0.047 | 0.025 | 0.062 |
| Day 3 ⇒ Cortisol Values | | 0.010 | 0.035 | 0.766 |
| Day 4 ⇒ Cortisol Values | | 0.046 | 0.037 | 0.213 |
| Day 5 ⇒ Cortisol Values | | 0.063 | 0.045 | 0.158 |
| Quadratic Term ⇒ Cortisol Values | | 0.009 | 0.002 | <0.001 |
| Sample-Specific Disturbances | A | 0.271 | 0.028 | <0.001 |
| | C | 0.000 | 0.000 | 0.431 |
| | E | 0.471 | 0.023 | <0.001 |
| *Level 2: Between-Person Variance Decomposition* | | | | |
| Hair Cortisol | A | 0.512 | 0.246 | 0.038 |
| | C | 0.711 | 0.154 | <0.001 |
| | E | 0.814 | 0.072 | <0.001 |
| Hair Cortisol ⇒ Waking Intercept | A | 0.152 | 0.143 | 0.287 |
| | C | -0.052 | 0.122 | 0.671 |
| | E | 0.019 | 0.036 | 0.608 |
| Waking Intercept Unique of Hair Cortisol | A | 0.275 | 0.107 | 0.011 |
| | C | 0.461 | 0.084 | <0.001 |
| | E | 0.268 | 0.040 | <0.001 |
| Hair Cortisol ⇒ Awakening Response | A | -0.337 | 0.308 | 0.274 |
| | C | 0.226 | 0.191 | 0.239 |
| | E | 0.096 | 0.092 | 0.300 |
| Waking Intercept Unique of Hair Cortisol ⇒ Awakening Response | A | -0.319 | 0.295 | 0.280 |
| | C | -0.103 | 0.244 | 0.674 |
| | E | -0.383 | 0.113 | 0.001 |
| Awakening Response Unique of Hair Cortisol and Waking Intercept | A | 0.410 | 0.213 | 0.055 |
| | C | 0.243 | 0.429 | 0.571 |
| | E | 0.343 | 0.070 | <0.001 |
| Hair Cortisol ⇒ Diurnal Slope | A | 0.019 | 0.012 | 0.111 |
| | C | 0.004 | 0.008 | 0.615 |
| | E | -0.001 | 0.003 | 0.766 |
| Waking Intercept Unique of Hair Cortisol ⇒ Diurnal Slope | A | -0.020 | 0.013 | 0.144 |
| | C | 0.000 | 0.008 | 0.992 |
| | E | 0.000 | 0.005 | 0.978 |
| Awakening Response Unique of Hair Cortisol and Waking Intercept ⇒ Diurnal Slope | A | 0.011 | 0.018 | 0.538 |
| | C | -0.004 | 0.028 | 0.900 |
| | E | 0.004 | 0.005 | 0.429 |
| Diurnal Slope Unique of Hair Cortisol Waking Intercept, and Awakening Response | A | 0.000 | 0.000 | 0.842 |
| | C | 0.034 | 0.007 | <0.001 |
| | E | 0.019 | 0.006 | 0.001 |
| *Covariates* | | | | |
| Age ⇒ Hair Cortisol | | -0.079 | 0.058 | 0.175 |
| Age ⇒ Waking Intercept | | -0.047 | 0.033 | 0.157 |
| Age ⇒ Awakening Response | | -0.006 | 0.053 | 0.915 |
| Age ⇒ Diurnal Slope | | 0.001 | 0.003 | 0.777 |

| | | | |
|---|---|---|---|
| Sex ⇒ Hair Cortisol | 0.407 | 0.106 | <0.001 |
| Sex ⇒ Waking Intercept | -0.099 | 0.042 | 0.017 |
| Sex ⇒ Awakening Response | -0.157 | 0.089 | 0.079 |
| Sex ⇒ Diurnal Slope | -0.003 | 0.005 | 0.590 |
| Age x Sex ⇒ Hair Cortisol | 0.006 | 0.108 | 0.958 |
| Age x Sex ⇒ Waking Intercept | 0.001 | 0.003 | 0.777 |
| Age x Sex ⇒ Awakening Response | 0.085 | 0.066 | 0.198 |
| Age x Sex ⇒ Diurnal Slope | 0.002 | 0.005 | 0.635 |
| *Conditional Means* | | | |
| Hair Cortisol | -0.050 | 0.060 | 0.410 |
| Waking Intercept | 1.627 | 0.045 | <0.001 |
| Awakening Response | 0.780 | 0.060 | <0.001 |
| Diurnal Slope | -0.342 | 0.021 | <0.001 |
| **Model Fit Indices:** -2Log Likelihood = -10868.946, AIC = 21843.891 | | | |

All estimates are unstandardized. Units are in raw concentrations residualized for batch year and log-transformed ⇒ Cholesky path. Age was standardized; sex was effect coded.
